# Evolution of dispersal syndrome and its corresponding metabolomic changes

**DOI:** 10.1101/178715

**Authors:** Sudipta Tung, Abhishek Mishra, Navdeep Gogna, Mohammed Aamir Sadiq, P.M. Shreenidhi, V.R. Shree Sruti, Kavita Dorai, Sutirth Dey

## Abstract

Dispersal is one of the strategies for organisms to deal with climate change and habitat degradation. Therefore, investigating the effects of dispersal evolution on natural populations is of considerable interest to ecologists and conservation biologists. Although it is known that dispersal itself can evolve due to selection, the behavioral, life-history and metabolic consequences of dispersal evolution are not well understood. Here we explore these issues by subjecting four outbred laboratory populations of *Drosophila melanogaster* to selection for increased dispersal. The dispersal-selected populations had similar values of body size, fecundity and longevity as the non-selected lines (controls), but evolved significantly greater locomotor activity, exploratory tendency, and aggression. Untargeted metabolomic fingerprinting through NMR spectroscopy suggested that the selected flies evolved elevated cellular respiration characterized by greater amounts of glucose, AMP and NAD. Concurrent evolution of higher level of Octopamine and other neurotransmitters indicate a possible mechanism for the behavioural changes in the selected lines. We discuss the generalizability of our findings in the context of observations from natural populations. To the best of our knowledge, this is the first report of the evolution of metabolome due to selection for dispersal and its connection to dispersal syndrome evolution.

## 1. Introduction

The biodiversity on earth is threatened today due to factors like rapid climate change, and habitat degradation and destruction (Root et al. 2003). These environmental stresses not only change the life-history and behaviours of species worldwide, but also affect their abundance and distribution patterns (Parmesan et al. 1999; Travis 2003). One of the primary strategies for mobile organisms to cope with such environmental challenges is to disperse to a more favourable habitat (Travis et al. 2013). Under such circumstances, evolution of greater dispersal could prove advantageous and play a crucial role in the persistence of populations. Furthermore, dispersal is known to be linked with several other ecological and evolutionary processes, such as, population stability (Dey and Joshi 2006), local adaptation (Gandon et al. 1996; Lenormand 2002), speciation (reviewed in Barton 2001), cooperation and sociality (Le Galliard et al. 2005), community dynamics (reviewed in Leibold et al. 2004), species invasion (Shaw and Kokko 2015), range expansion (Travis and Dytham 2002) and disease spread (Rappole et al. 2006). Thus dispersal evolution can be expected to have pronounced effects at the population and community level (Brown and Kodric-Brown 1977; Leibold et al. 2004). Not surprisingly, the causes and consequences of dispersal evolution have been a major focus of investigation for the past decade (Fronhofer and Altermatt 2015; Williams et al. 2016; Fronhofer et al. 2017; Ochocki and Miller 2017; Weiss-Lehman et al. 2017).

Theoretical and empirical studies have shown that dispersal can evolve in response to several ecological scenarios, e.g., spatially correlated extinctions (Fronhofer et al. 2014), kin competition (Gandon 1999; Billiard and Lenormand 2005) and inbreeding depression (Guillaume and Perrin 2009). However, it is rather difficult to make generic predictions about the effects of dispersal evolution on other life-history and behavioural traits. This is because, *inter alia*, dispersal is a composite trait, consisting of three stages, namely emigration from the natal habitat, inter-patch movement and immigration into the destination patch (Bowler and Benton 2005). The kind of environmental and the corresponding physiological and behavioural challenges faced during each of these stages can be very different. As a result, when under selection for dispersal, evolution of a particular trait critically depends on how that trait influences the different stages of dispersal, the nature of the selection pressure faced in each stage, the costs associated with them, how these costs interact with each other, how they are countered by the organisms and the underlying genetic constraints for the evolution of the focal trait. Evidently, this makes *a priori* prediction of the correlated response of dispersal selection rather difficult.

Among the life-history traits, the most likely candidate to evolve in response to selection for greater dispersal is body size. This is because as dispersal is an energy-intensive trait (Bonte et al. 2012; Travis et al. 2012) bigger individuals should have more resources and therefore be more successful dispersers. Consistent with this notion, body size is found to be positively associated with dispersal in several taxa, *inter alia*, milkweed bugs (Dingle et al. 1980), butterflies (Sekar 2012), birds (Dawideit et al. 2009) and mammals (Sutherland et al. 2000). Does this mean that when dispersal evolves as a trait, body size will co-evolve? Although this is one of the most well-established trait associations in the dispersal literature, it is also known that the existence of a correlation between two-traits does not guaranty that they will co-evolve (Leroi et al. 1994). Unfortunately, to the best of our knowledge, the body size-dispersal correlation has never been demonstrated in an evolutionary context.

Two other major life-history traits that have been shown to have an association with dispersal are fecundity and longevity. This is thought to be intuitive, since, in order to accommodate the high energy demands of dispersal, the normal resource allocation patterns of the organisms are expected to alter (Bonte et al. 2012). But in reality, the relationship between these traits is not straightforward. In wing-dimorphic insects (reviewed in Roff 1977) and soil nematode *Caenorhabditis elegans* (Friedenberg 2003), dispersers are found to have lower fecundity, whereas the opposite is observed in field voles *Microtus agrestis* (Ebenhard 1990) and no difference is found in parasitoid wasps (Innocent et al. 2010). Similarly, although positive covariation between dispersal and lifespan has been theoretically predicted in the context of kin competition (Dytham and Travis 2006), laboratory experimental evolution studies have reported that these two traits do not coevolve (Fronhofer et al. 2014) as a correlated response to selection for dispersal.

The other class of traits that have been extensively investigated in the context of dispersal are those related to behaviour. For example, locomotor activity is found to covary positively with dispersal in both single-generation trait-association studies (in butterfly *Melitaea cinxia*, Hanski et al. 2006) as well as artificial selection experiments in red flour beetle *Tribolium castaneum*, (Matsumura et al. 2016). Similarly, efficient exploration of the nearby areas is expected to be beneficial for successful dispersers and exploratory tendency is often found to be strongly associated with dispersal (Korsten et al. 2013). Interestingly, in many species, exploration is also found to be strongly related to invasion (Rehage et al. 2005; Cote et al. 2010; Russell et al. 2010), which involves conflict with the native species. Thus, not surprisingly, aggression is another behavioural trait strongly correlated with exploration (Verbeek et al. 1996; Dingemanse and de Goede 2004) and is often closely related to personality-dependent dispersal (Cote et al. 2010). These traits, which have a consistent association with dispersal, are often collectively termed as a part of ‘dispersal syndrome’ (Clobert et al. 2009). Although the notion of dispersal syndrome is well established in the literature (Ronce and Clobert 2012), its concurrent evolution with dispersal has been rarely explored. Moreover, it is difficult to comprehend whether the enhanced expression of the syndrome traits in natural populations is due to an independent advantage of those traits in the specific ecological scenarios or as a consequence of their genetic correlation with dispersal (Ronce and Clobert 2012). In order to investigate this problem, one needs to study dispersal evolution in an environment devoid of any apparent parallel selection pressure acting on the syndrome traits, which is difficult to attain under natural settings.

Another aspect of dispersal that has received some attention is the molecular basis of the dispersive phenotype (for a review see Saastamoinen et al. 2017). Unfortunately, the polygenic nature of most dispersal traits (Zera and Brisson 2012) and the inherent complexity of the dispersal process itself (Bowler and Benton 2005), indicates that there are relatively few single genes that have been shown to be associated with dispersal. Two such genes in insects are the glycolytic enzyme encoded by Phosphoglucose isomerase gene *(pgi*) in the Glanville fritillary butterflies (Niitepõld et al. 2009) and the cGMP-dependent protein kinase called foraging *(for*) gene in *Drosophila melanogaster* (Osborne et al. 1997). Interestingly, although the *for* gene was discovered in the context of larval locomotion (de Belle et al. 1989), it has also been shown to play a role in adult locomotion and dispersal (Pereira and Sokolowski 1993; Edelsparre et al. 2014). In *C. elegans* as well, three genes, namely G-protein coupled receptors *npr-1* (de Bono and Bargmann 1998), *tyra-3* (Bendesky et al. 2011) and *rol-1* (Friedenberg 2003), have been shown to be associated with dispersal phenotypes. However, as alluded earlier, the polygenic nature of many dispersal traits makes it impossible to predict whether these are the genes whose frequencies would change to accommodate the excess energy demand of dispersal and associated behavioural changes during dispersal evolution. A different and more fruitful approach to understand the sub-cellular basis of dispersal evolution perhaps would be to investigate the metabolome of the dispersers and try to elucidate the various processes that have been up-or down-regulated (Matsumura et al. 2016). For example, it is known that a number of neurotransmitters - octopamine, dopamine, serotonin and tyramine - play a crucial role in determining several behaviours in insects, including activity, exploration and aggression (Yellman et al. 1997; Zhou et al. 2008), all of which are known to be associated with dispersal (Wahlström 1994; Hanski et al. 2006; Duckworth and Badyaev 2007; Cote et al. 2010; Korsten et al. 2013). Unfortunately, the metabolomic changes accompanying dispersal evolution remain unexplored.

In order to address some of the issues mentioned above, we investigated the correlated response to selection for increased dispersal in four large, outbred populations of *Drosophila melanogaster.* These populations have been subjected to artificial selection for the longest duration (69 generations, ongoing) documented in the dispersal literature, using a setup that mimics increasing habitat fragmentation over generations (Tung et al. 2018). The selected populations have rapidly evolved significantly greater dispersal propensity and ability (i.e. distance covered by the dispersers) and a larger fraction of Long-Distance Dispersers (LDDs) in the population (Tung et al. 2018). Here, we assayed three behavioural traits (locomotor activity, exploration and aggression) and three life-history traits (body size, fecundity and life-span) of the selected lines and contrasted the results with corresponding controls. We found that the behavioural traits had evolved while the life history traits had not. Further, we compared the metabolome of the selected and the control flies using non-targeted NMR spectroscopy and connected them with the corresponding behavioural changes.

## 2. Materials and methods

The details of all assays are mentioned in the Appendix S1 in Supporting Information (Text S1) and only a brief description is presented here as an aid to appreciate the results and discussion.

### 2.1 Ancestry and maintenance regime of the experimental populations

We used eight large (breeding size of ~2400) laboratory populations of *Drosophila melanogaster*, four of which (VB_1-4_, VB stands for ‘**V**aga**b**ond’) had been selected for greater dispersal ability and propensity for 67 generations (when the last set of assays reported here was performed) and the remaining four (VBC_1-4_, VBC stands for ‘**VB-C**ontrol’) were their corresponding ancestry-matched controls (Tung et al. 2018). The VB_1-4_ and VBC_1-4_ flies were derived from the corresponding DB_1-4_ populations (Sah et al. 2013). Thus, VB_i_ and VBC**j** populations that share a subscript (i.e. *i* = *j*) are related by ancestry, and hence were always assayed together and treated as blocks in statistical analyses. The ancestry of these flies traces back to the IV lines, which were wild-caught at South Amherst, MA, USA in 1970 (Ives 1970). These flies have been maintained ever since at large population sizes (to ameliorate inbreeding-like effects) under laboratory conditions.

Both VB and VBC populations were maintained on a 15-day discrete generation cycle. For each population in each generation, eggs were collected in clear ~37-mL plastic vials at a density of 60-80 eggs per ~6 mL of standard banana-jaggery medium (following Sheeba et al. 1998). On the 12^**th**^ day after egg collection, the adults from VB populations were subjected to selection for dispersal (see the following paragraph for details). After this, the adults were transferred to a plexi-glass cage (25 cm ×20 cm ×15 cm) and supplied with live yeast-paste to boost fecundity. On the 14^th^ day after egg-collection, the adults were allowed to lay eggs on a petri-plate containing banana-jaggery medium. These eggs were then dispensed into plastic vials to form the next generation, while the adults were discarded. The breeding adult population of both VBs and VBCs were ~2400 individuals. The details of the maintenance of these populations can be found as Text S1.1.

### 2.2 Selection protocol

The apparatus used for dispersal selection had three components- *a source, a path and a destination* (Tung et al. 2018). In order to impose selection in a given VB population, the adults collected from 80 vials (~4800 individuals) were divided equally in two sets and introduced into the *sources* (plastic containers of volume ~1.5 L) of two independent dispersal set-ups. Each *source* was then connected to the *destination* (a plastic container similar to the source) using the *path*, which was a plastic tube of inner diameter ~1 cm.

Length of the path was increased over generations (see below). The flies dispersed from the source, through the *path*, into the *destination*. The source and the path did not contain any food or moisture, while the destination contained a strip of moist cotton. The flies were allowed to disperse until ~50% of the population reached the *destination* (estimated visually) or for six hours, whichever happened earlier. The duration of six hours was chosen because prior studies in the lab showed that the flies did not experience any death due to desiccation during this period. Only the flies which reached the *destination* in the two selection setups were transferred into a cage and allowed to breed for the next generation. Since only 50% of the population in a given dispersal set-up reached the destination, we used two such set-ups for each VB population. This ensured that the breeding population sizes of the selected and the control flies (i.e. the VBCs) were similar.

In each VBC population, adults from 40 vials were introduced into a *source* container, but the opening of the container was plugged so that the flies could not disperse. They were also provided with a moist cotton plug, after 25% of the VBs reached their destination or 3 hours (whichever was earlier). Thus the VBCs also experienced similar desiccation stress as the VBs, and therefore, the observed differences between VBs and VBCs could be attributed to the selection for dispersal. All the flies from a given VBC population were transferred to the cages and allowed to breed, thus ensuring that there was no selection for dispersal.

The length of the path was 2 m at the beginning of the experiment but was increased intermittently over generations in order to intensify the selection for grater dispersal ability. By generation 67, when the last set of assays was performed, the path length had reached 20 m.

### 2.3 Assays

All assays were performed after rearing both VB and VBC populations for one generation under common conditions to minimize the contributions of non-genetic parental effects (Watson and Hoffmann 1996). For all assays, we collected eggs from the VBs or the VBCs on the 15^th^ day after egg-collection and reared them under identical conditions (60-80 eggs per ~6 mL food). The adult flies thus obtained were maintained identically for VBs and VBCs (25°C, constant light conditions) in plexi-glass cages and served as parents for the flies which were used in various assays. While collecting eggs for the assays, for both VBs and VBCs, a low egg density (~50 eggs on ~6mL food) was maintained to minimize the effects of larval crowding on the life history and behavioural traits measured. The environmental (constant light, 25°C temperature, 80-90% humidity, abundant nutritional availability, low rearing density) and physiological (similar age) conditions were controlled and maintained identically across all VB and VBC populations for all the following assays.

#### 2.3.1 Behavioural assays

Dispersal is often found to be positively correlated with a number of behavioural traits like locomotor activity, exploration and aggression (Hanski et al. 2006; Duckworth and Badyaev 2007; Cote et al. 2010; Korsten et al. 2013; Matsumura et al. 2016). Thus, in order to investigate the correlated response to dispersal evolution, we compared these traits between the selected and control lines.

Locomotor activity and resting behaviour of adult male flies of the VB and VBC populations were assayed using the *Drosophila* Activity Monitoring (DAM2) data collection system (Trikinetics Inc, Waltham, MA) following standard protocols (Chiu et al. 2010; see also Text S1.2.1). During assay, individual flies were introduced into small (length: 6.5 cm, diameter: 5 mm) glass tubes. These tubes were placed in the monitoring apparatus such that two independent perpendicular IR beams passed through the centre of each glass tube. The activity for a given fly was estimated as the average number of times the fly crossed the IR beam per hour (Chadov et al. 2015). Continuous inactivity for five minutes or more was considered as sleep/rest (Hendricks et al. 2000; Chiu et al. 2010). For each VB/VBC population, we recorded two kinds of activity/rest data on 30-32 12-day old (post egg-collection) male flies. Prior studies on several insects, including *Drosophila*, have reported that when the organisms are introduced into a novel environment, activity levels are greater than their normal level of spontaneous activity (Liu et al. 2007). To capture this initial hyperactivity in the selected and control flies in the presence and absence of food (Tung et al. 2018), we recorded the activity for the first 6 h after introducing the flies into the glass tubes. This duration was the same as what was given to the flies to disperse during the selection process. We also recorded the basal level of spontaneous activity of these flies in presence of food over a 24-h duration after 6 h of acclimation. For each of these three datasets, average activity counts per hour and the fraction of time spent in sleeping/resting were computed. Note that activity and rest are defined on different stretches of time, and therefore are independent of each other. For example, a given fly can potentially rest/sleep for longer period and still be more active by running back-and-forth excessively in the tube, while it is awake/not resting. In principle, unequal levels of rest/sleep interruptions can lead to differences in sleep-related stress levels, even across organisms whose total duration of sleep is the same. In order to investigate this possibility, we compared the quality of rest/sleep of the selected and control lines by measuring the average length of uninterrupted rest/sleep bout and duration of the longest rest/sleep bout for each of the flies from the 24-h dataset (Chiu et al. 2010).

Exploratory tendency was measured on the 12^th^ day post-egg collection, following an earlier assay setup (Liu et al. 2007), using 32 flies of each sex from each VB and VBC population. For this assay, the flies were introduced singly into an enclosure formed by placing a petri-plate lid (10 cm diameter) on a white background (see Text S1.2.2 for details). In such a setup, the flies generally prefer to walk along the boundary of the enclosure (i.e. side-wall of the lid), avoiding venturing closer to the centre (Soibam et al. 2012). Therefore, the number of times a fly goes more than an arbitrary distance away from the boundary can be taken as a measure of its exploratory tendency. Here, following previous studies (Liu et al. 2007; Soibam et al. 2012), this distance was taken to be ~1 cm (see Text S1.2.2 for details).

Male-male aggression was estimated from 30 VB-VBC fights for each block (i.e. VB_1_-VBC_1_ and so on), based on existing protocols in the literature (Yurkovic et al. 2006; see Text S1.2.3 for details). Briefly, freshly eclosed males were collected and reared in social isolation (i.e. one male per vial) till the day of assay. On the 12^th^ day post-egg collection, two males (one VB and one VBC) were introduced into an aggression setup and their interaction was recorded for 45 minutes using a video camera. The males were colour-coated with daylight fluorescent pigments (DayGlo) for easy identification (see Text S1.2.3 for details). The setup consisted of one well of a 12-well culture plate (Corning®, NY, USA), with a small plastic cup containing regular banana-jaggery medium affixed at its centre. A freshly decapitated female was stuck to the centre of this food cup using yeast paste. The food cup served as the aggression arena. Following a previous study (Yurkovic et al. 2006), a winner in a fight was declared when one male chased the other male off the arena three consecutive times. For each fight, the identity of the winner *(i.e.* VB or VBC) was recorded and the total number of wins by the two groups were compared.

#### 2.3.2 Life-history assays

In terms of life-history, we investigated body size (measured as dry weight), fecundity and longevity: three traits that were likely to have evolved in the selected lines in order to accommodate the excess energy need for dispersal.

Dry weight was estimated on five batches of 20 males or 20 females from each of the four VB and VBC populations, after drying at 60°C for 72 h (for details see Text S1.2.4).

For fecundity assays, the number of eggs laid within a 12-h period by 15-day old or 33-day old females was estimated as early-life and late-life fecundity respectively (Text S1.2.5). For each population × reproductive timing combination, we used 40 pairs (1 male + 1 female) of flies for estimating fecundity.

For the longevity assay, we monitored 10 sets of 10 flies of either sex per population daily, from eclosion till death. Longevity was scored as the number of days to death for each fly (Text S1.2.6).

#### 2.3.3 NMR sample preparation and spectroscopy

As dispersal is an energy-intensive process, in order to maintain an elevated level of dispersal phenotype and to cope with its high energy demand (Bonte et al. 2012), metabolism of the selected flies is expected to undergo considerable changes. To investigate this, we performed an untargeted metabolomics study using NMR spectroscopy on one block of selected-control populations (VB_4_-VBC_4_). This study allowed an untargeted analysis of the metabolome of the two populations and a comparison of the relative levels of different metabolites in the selected and control populations. For each population, we used 11 replicates, each of which consisted of 30 males. Samples were prepared following established protocols (Gogna et al. 2015) and subjected to both 1D and 2D NMR experiments. This was followed by metabolite fingerprinting using databases such as MMCD (http://mmcd.nmrfam.wise.edu) and BMRB (http://www.bmrb.wise.edu) (for details see Text S1.2.7).

### 2.4 Statistical analyses

VB_i_ and VBC_j_, that shared a subscript (i.e. *i* = *j*) were assayed and analysed together as a block as they were related to each other by ancestry. For locomotor activity, the fraction of time spent resting/sleeping (arcsine-square root transformed), average sleep bout, length of the longest sleep bout and female fecundity data, we used two factor mixed-model ANOVA with selection (VB and VBC) as a fixed factor crossed with block (1-4) as a random factor. Three-factor mixed-model ANOVA was performed for adult exploration, dry body weight and mean life span, with selection (VB and VBC) and sex (male and female) as fixed factors and block (1-4) as a random factor, crossed with each other. For aggression, the number of wins for VBs and VBCs were computed for all four blocks and the difference between the VB-VBC was tested using Wilcoxon Matched Pairs Test. We also computed the effect size (Cohen’s *d)* for all the above differences between the means. The value of *d* was interpreted as large, medium and small for *d*≥0.8, 0.8>*d*≥0.5 and *d*<0.5, respectively (Cohen 1988). All the above statistical analyses were performed using STATISTICA^®^ v5 (StatSoft. Inc., Tulsa, Oklahoma).

The longevity data were further analysed to obtain Irwin’s restricted mean lifespan (Irwin 1949) from the Kaplan-Meier survivorship estimates (Kaplan and Meier 1958) for all the VB and VBC populations. The statistical analysis for Irwin’s restricted mean lifespan was similar to the analysis of the mean lifespan. We also analysed the longevity profiles of these populations using Cox proportional hazards for two different models. In the mixed effects model ‘Block’ (4 levels, 1-4) was the random factor crossed with selection (VB/VBC) and sex (male/female), which were the fixed factors. We used the ‘coxme’ (version 2.2-5) package (Therneau 2015a) on R 3.4.0 (R Core Team 2017) for this analysis. We also used a Cox proportional hazards regression model without including Block as a factor, using the coxph function of the ‘Survival’ (version 2.41-3 package (Therneau 2015b) in R. For both analyses, the flies lost during the assay (overall less than 4%) were included as ‘right’ censored (zero censored) data.

The NMR spectral data were analysed using standard procedures (see Text S 1.2.7). The spectral data were normalized to total area, Pareto-scaled and subjected to Principal Component Analysis to identify and remove the outliers. This was followed by Orthogonal Projections to Latent Structure-Discriminant Analysis (OPLS-DA) to identify the metabolites responsible for separating VB and VBC flies. The significance test of the model was performed using CV-ANOVA (cross-validated ANOVA). Further, permutation analysis was performed on the best model using 1000 permutation tests with a threshold *P*-value of 0.05, which indicated that less than 5% of the results were better than the original one. The average level of the metabolites in the selected and control populations were compared using Student’s *t*-tests, followed by Bonferroni correction, thus restricting the family-wise error rate to <0.05.

## 3. Results

### 3.1 VBs were restless but rested less

During the first 6 h after set-up, the VB populations had significantly greater locomotor activity irrespective of the absence (Fig. 1A, F_1,3_=60.3, 0.004) or presence (Fig. 1C, F_1,3_= 423.3, *P*=0.0003) of food. Moreover, of the total duration of 6 h, the VBs spent significantly less time in rest/sleep, both in the absence (Fig. 1B, F_1,3_=50.4, *P*=0.006) and presence (Fig. 1D, F_1,3_=386.9, *P*=0.0003) of food. Interestingly though, when assayed in the presence of food over a 24-h duration after the initial 6 h of acclimatization, although the difference in activity persisted (Fig. 1E, F_1,3_= 59.9, *P*= 0.004), the VBs spent similar amount of time in rest/sleep as the VBCs (Fig. 1F, F_1,3_= 5.47, *P*=0.1). Over 24 h, the length of average sleep bouts (Fig. S1A, F_1,3_=2.2, *P*=0.23) and maximum sleep bouts (Fig. S1B, F_1,3_=4.7, *P*=0.12) of the VBs and the VBCs were also comparable.

**Figure 1.**
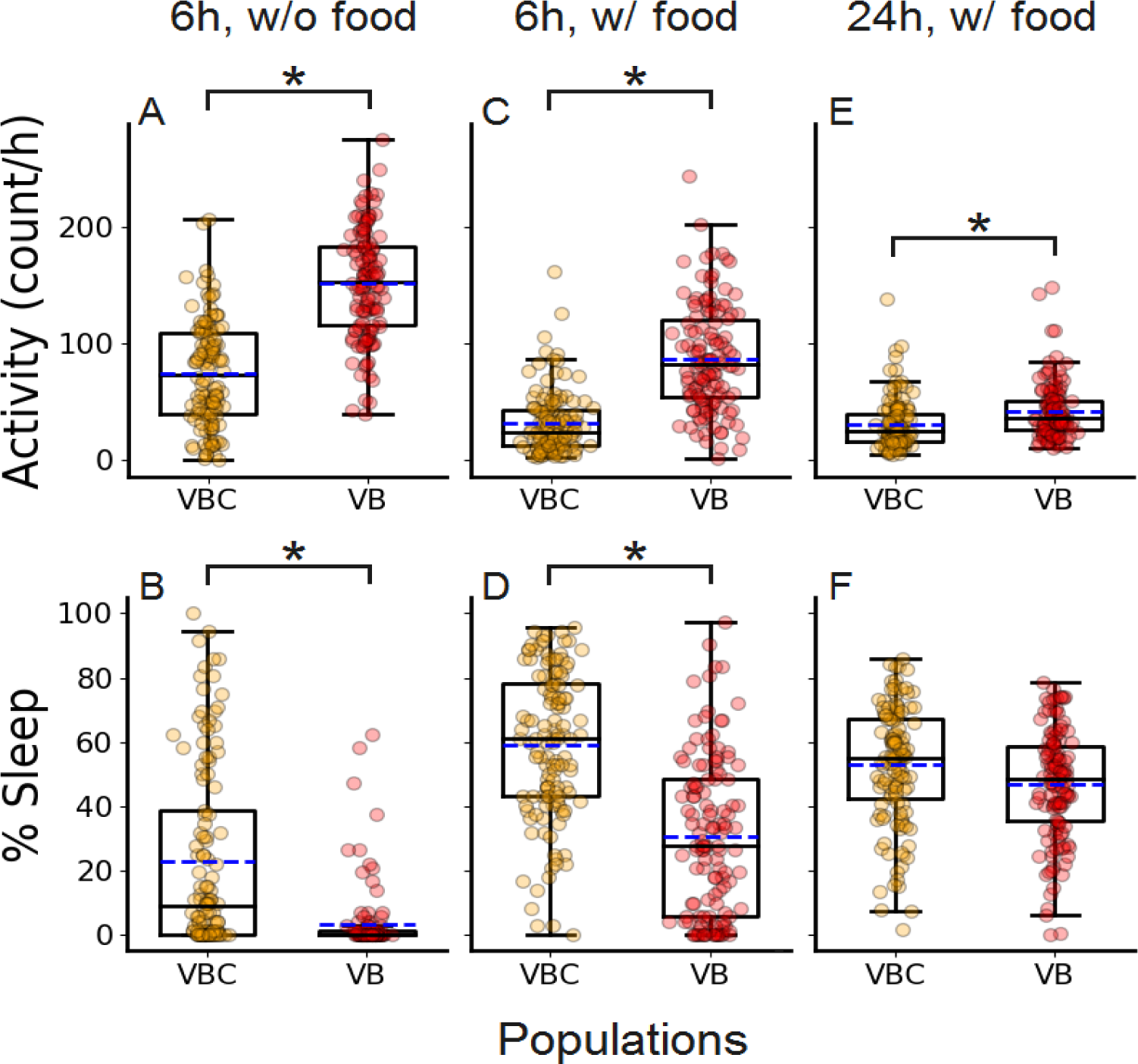
Locomotor activity-sleep profiles in the presence and absence of food. Cleveland-box plots show that in the absence of food during the first 6 h after introduction, **(A)** locomotor activity of VBs was significantly greater than the VBCs, although **(B)** duration of rest was significantly lower. In the presence of food during the first 6 h, similar results were obtained for **(C)** locomotor activity and **(D)** rest. After acclimatization for 6 h, over the next 24 h, the VBs had significantly greater (**E**) locomotor activity but similar levels of (**F**) sleep duration as the VBCs. The points represent the pooled data for all the replicates of VB and VBC populations, with small random jitter along the X-axis. The edges of the box denote 25^th^ and 75^th^ percentiles, while the black solid lines and blue broken lines represent the median and mean respectively. The error bars represent standard errors around the mean (SEM) and * denotes *P*<0.05.

### 3.2 Selection for dispersal led to greater exploratory behaviour and aggression

Flies from VB lines made more exploratory trips than those from VBC lines when introduced into a novel environment (F_1, 3_=11.96, *P*=0.04). In general, males showed greater exploratory behaviour than females (F_1, 3_=27.12, *P*=0.01), but no sex × selection interaction was found (Fig. 2A, F_1, 3_=0.009, *P*=0.93). The VB males were more aggressive than the VBC males (Wilcoxon Matched Pairs Z= 1.83, *P* = 0.068) and the effect size of this difference was very large (*d* = 2.05) (Fig. 2B).

**Figure 2.**
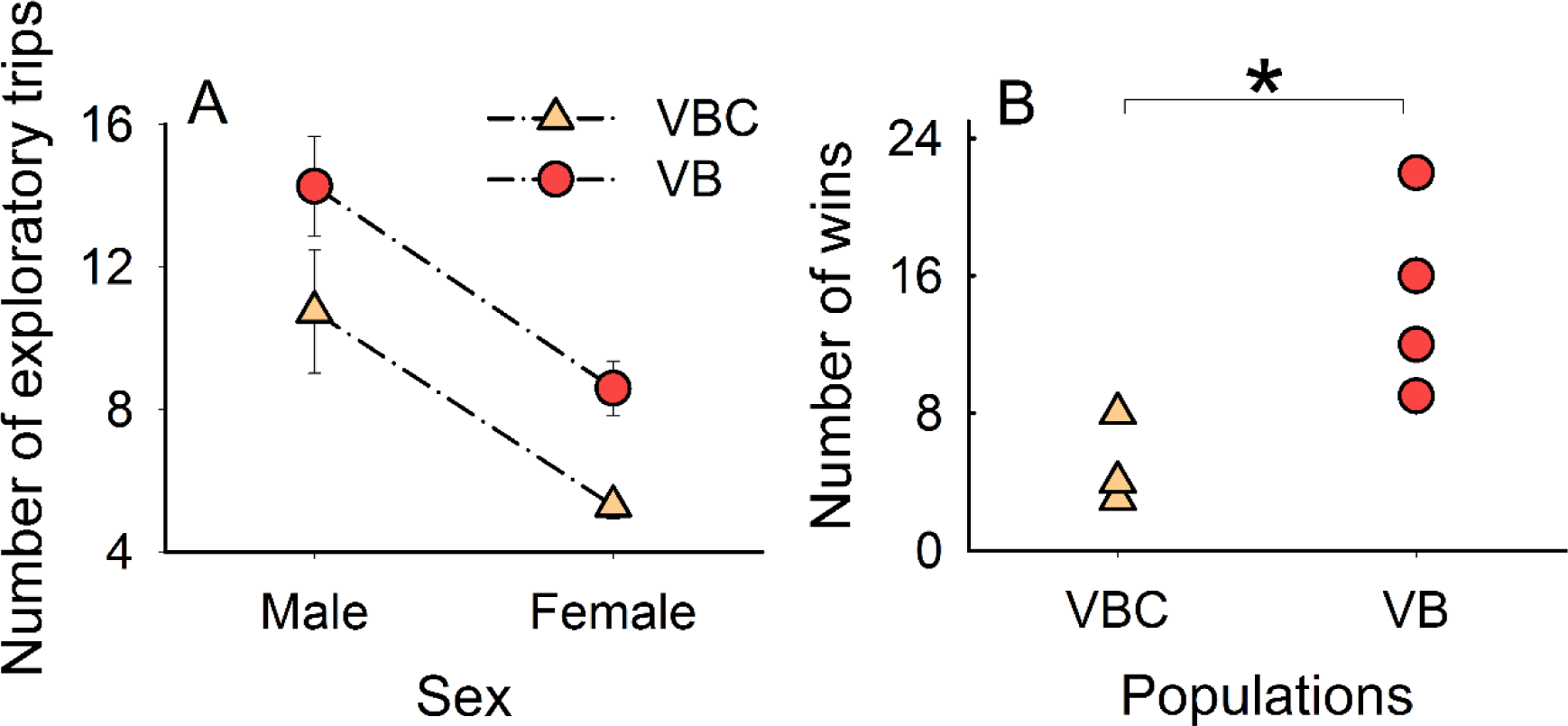
Exploration and aggression behaviour of VBs and VBCs. (**A**) VB males and females undertook significantly greater number of exploratory trips than VBCs. The error bars represent standard errors around the mean (SEM). (**B**) VB males were more aggressive as they won significantly higher number of fights against VBC males. Both these results were consistent across all the four blocks. * denotes *P*= 0.068, with a very large effect size (Cohen’s *d*= 2.05).

### 3.3 Selection for dispersal did not lead to changes in dry body weight, fecundity or longevity

The mean dry weights of VBs and VBCs were found to be comparable (Fig. 3A, F_1, 3_= 0.76, *P*=0.45). Although *Drosophila* females are known to be heavier than males, we did not find any significant selection × sex interaction (F_1, 3_= 2.1, *P*=0.24) in our study, suggesting that selection did not affect the dry weight of the two sexes in VB and VBC flies differentially. There was no significant difference between the fecundity of the VB and VBC flies with respect to either early fecundity (Fig. 3B, F_1, 3_= 0.25, *P*=0.65) or late fecundity (Fig. 3B, F_1, 3_= 0.2, *P*= 0.68), indicating the absence of a trade-off between increased dispersal ability and reproductive output. We did not find any trade-off between dispersal and longevity either: the average life-span of VBs was similar to that of the VBCs (Fig. 3C, F_1, 3_= 4.9, *P*= 0.11). There was no difference between the VBs and the VBCs in terms of Irwin’s restricted mean lifespan, computed from the corresponding Kaplan-Meier estimates (Fig. S2, F_1, 3_= 5.88, *P*= 0.094). Overall survivorship profiles of VBs and VBCs were also found to be similar, when they were compared using Cox proportional hazards model including the random factor (z = - 45 *P* = 0.65) as well as excluding the random factor (z = -1.24, *P*= 0.214).

**Figure 3.**
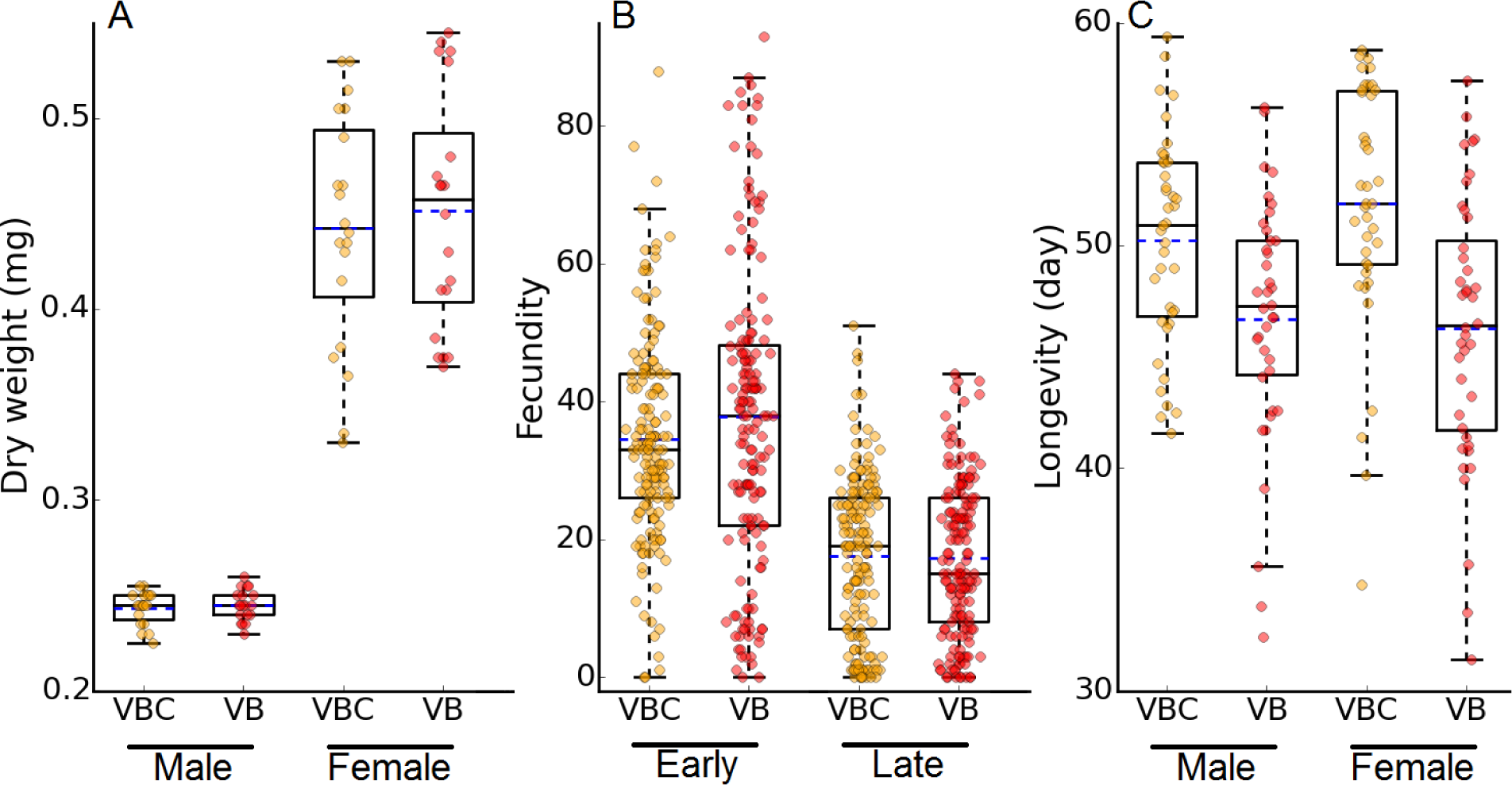
Life-history traits of VBs and VBCs. Cleveland-box plots show (**A**) dry body weight, (**B**) early-late fecundity and (**C**) longevity profiles of VB and VBC populations, which were not different from each other. The points represent the pooled data for all the replicates of VB and VBC populations, with small random jitter along the X-axis. The edges of the box denote 25^th^ and 75^th^ percentiles, while the black solid lines and blue broken lines represent the median and mean respectively.

### 3.4 Selection for dispersal led to changes in the metabolome profile

OPLS-DA scores plot (Fig. S3) documents the differences in the metabolite profile of the selected and control flies. Fig. S4 shows the colour-coded coefficient loadings plot used to identify the metabolites responsible for differentiating VB and VBC flies. The variance explained by the model *(R*^*2*^ *X)* was 0.968 and the variance predicted by the model (*Q*^*2*^) was 0.953, showing that the model was effective and had a good predictive accuracy. The credibility and robust nature of the model were also confirmed by testing the statistical significance of the model with CV-ANOVA (p-value <0.01) and permutation test (p-value <0.05). The metabolites, which were significantly different between the selected and control flies in *t*-test followed by Bonferroni correction, are tabulated in Table 1. The functional importance of these metabolites in the context of dispersal evolution has been discussed in sections 4.6-4.8.

**Table 1:** List of significantly altered metabolites during the course of selection for increased dispersal.

| **Metabolites** | ***p* values**<br>**after**<br>**correction** | **Effect size**<br>**(Cohen's *d*)** | **Fold change in VB**<br>**(VB/VBC)** |
| --- | --- | --- | --- |
| Tryptophan | 1.29E-04 | 3.77 | 3.27 |
| AMP | 7.76E-04 | 3.16 | 2.63 |
| 3-HK | 1.07E-04 | 3.86 | 2.5 |
| Octopamine | 6.51E-06 | 3.88 | 2.26 |
| Phenylalanine | 1.68E-04 | 3.58 | 1.95 |
| NAD | 4.52E-06 | 4.8 | 1.82 |
| Tyrosine | 6.93E-06 | 3.96 | 1.73 |
| Histidine | 3.05E-03 | 2.72 | 1.71 |
| Glucose | 2.70E-05 | 4.18 | 1.65 |
| ATP | 1.19E-04 | 3.69 | 1.57 |
| Sucrose | 2.4E-02 | 2.39 | 1.48 |
| Citric acid | 1.76E-02 | 2.15 | 1.36 |
| Lactic acid | 7.72E-03 | 2.26 | 1.35 |
| Fatty acid | 2.39E-02 | 2.49 | 0.77 |

Note: The *p*-values for a given metabolite were obtained from *t*-tests between the VBs and VBCs, followed by Bonferroni correction to restrict the familywise error rates. All effect size values are large (i.e. *d*>0.8). Note that the AMP: ATP ratio for the VBs and VBCs were 0.43 and 0.26 respectively.

## 4. Discussion

### 4.1 Selection for dispersal leads to similar patterns of activity but different patterns of rest in the short and long time-scales

During the first 6 h after introduction to a new environment (same duration for which the flies are allowed to disperse during selection), the selected populations had greater locomotor activity than the controls (Fig. 1A and 1C), and spent lesser time in rest (Fig. 1B and 1D). This observation is consistent with previous results (Hanski et al. 2006; Matsumura et al. and also with the fact that the VBs were under intense selection to reach a new environment within the first 6 h of introduction to the source (Tung et al. 2018). Consequently, maximizing the amount of activity and minimizing the resting period during that time would be of obvious advantage to the VB flies. More interestingly though, we found similar activity/rest patterns in the absence and presence of food *(cf* Fig. 1A with 1C and 1B with 1D), which suggests that the increased activity is independent of starvation or desiccation cues. This is again consistent with earlier observations that the dispersal propensity and ability differences between VBs and VBCs were observed irrespective of the presence or absence of food (Tung et al. 2018). However, when we measured the activity of these flies over 24 h, after allowing a period of 6 h to acclimatize, although the VB males were found to be significantly more active than VBC males (Fig. 1E), the percentage of time spent resting was not different between these two lines (Fig. 1F). In other words, the VB flies rest less during the period that corresponds to the time when they face selection, but revert to normal levels of rest once that phase is over. During the latter period, the quality of the rest/sleep of the flies, as measured by the length of average bout of sleep (Fig. S1A) and maximum bout of sleep (Fig. S1B), during 24 h, was also found to be similar in VBs and VBCs. Thus, although the VBs are more active, it seems unlikely that they would face negative effects of rest-deprivation in the long run. To the best of our knowledge, this is the first demonstration that dispersers also modulate their rest-patterns temporally in way that could reduce the negative effects of rest-deprivation (Huber et al. 2004; Kayser et al. 2015). Increased activity of dispersers can positively correlate with another important behavioural trait, namely exploratory tendency (Cote et al. 2010), which we investigated next.

### 4.2 The evolution of dispersal led to simultaneous evolution of exploratory behaviour

Dispersers often have greater exploratory tendency (Cote et al. 2010; Korsten et al. 2013), which is thought to be beneficial for finding new habitats. In our selection protocol, there was no sensory cue in the path connecting the source to the destination. Therefore, only those flies (of either sex) could disperse successfully that took the risk of getting into the path and then continuing along it, implying that exploratory tendency was under strong positive selection. Therefore, it was no surprise that the dispersal-selected VBs were more exploratory in nature than the VBCs, and the result was consistent across both sexes (Fig. 2A). An overall higher number of exploratory trips by the males than the females agrees with a previous observation that the latter sex is intrinsically more centrophobic in this species (Besson and Martin 2005). Elevated exploratory tendency in dispersers can be important during range expansion as the individuals present at the range edges are more likely to experience environments different from their native and/or previously introduced habitats. For example, Kenyan house sparrows that were present at a range expansion front were found to be significantly more exploratory (Liebl and Martin 2012). Interestingly, in many species, exploration is also found to be strongly related to invasion (Rehage et al. 2005; Cote et al. 2010; Russell et al. 2010), which involves conflict/confrontation with the native species. Consequently, aggression is another behavioural trait strongly correlated with exploration (Verbeek et al. 1996; Dingemanse and de Goede 2004) and often closely related to personality-dependent dispersal (Cote et al. 2010). Thus, we next investigated the effect of dispersal evolution on aggression.

### 4.3 Male-male aggression evolved as a correlated response to selection for dispersal

Aggression is an important trait that influences an individual’s ability to retain resources and mates or gain new ones (O’Riain et al. 1996). Not surprisingly, several studies have reported a strong association between dispersal tendencies and aggression (Wahlström 1994; Duckworth and Badyaev 2007) which is consistent with our observations (Fig. 2B). However, while enhanced aggression might be a factor for dispersal success in some of the natural populations (Duckworth and Badyaev 2007), in our system, the dispersing flies had no obvious fitness advantage for being more aggressive, as they did not have to compete with any native individuals at the destination. Thus, in our experiment, aggression has likely evolved as a correlated response of dispersal evolution. This conclusion was strengthened when we investigated the changes in the metabolite levels of the selected flies (see section 4.7 below).

### 4.4 Dispersal-selected lines have comparable body size as that of the controls

One life history trait that could potentially explain the increased levels of dispersal, locomotor activity and aggression in the VBs is adult body size. Bigger organisms are expected to have greater energy reserves and, in general, body size is positively correlated with dispersal (Dingle et al. 1980; Sutherland et al. 2000; although see Gu and Danthanarayana 1992). Moreover, in *Drosophila melanogaster*, it is known that larger males win significantly more aggressive encounters compared to smaller males (Partridge and Farquhar 1983). However, we failed to find a significant difference in body weight, a proxy for body size, between VBs and VBCs (Fig. 3A), which suggests that the increased aggression and locomotor activity in VBs were not mere artefacts of differences in body sizes between the two populations. This result also highlights that how a trait evolves (here, no change in body size) due to a particular selection pressure may not always be inferred from existing trait-associations (here, the general observation from literature that body size and dispersal are positively correlated).

In *Drosophila*, body size is generally considered to be a good proxy for the total amount of resources available to an organism. Our results suggest that the selected flies have similar levels of resources compared to the controls (Fig. 3A), but at the same time, display elevated levels of activity (Figs 1A, 1C and 1E). Given that the kind of nutrients available to both populations are the same, one way for the selected flies to manage this feat would be to alter the pattern of resource allocation among the various traits (van Noordwijk and de Jong 1986). To investigate this possibility, we measured two crucial life-history traits, namely fecundity and longevity.

### 4.5 Increased dispersal does not affect fecundity or longevity

The relationship between dispersal and fecundity has been somewhat controversial in the literature. On one hand, flight ability/dispersal has been shown to be negatively correlated with fecundity in several insects including *Drosophila* (Roff 1977), long-winged crickets (Roff and Fairbairn 2007) and aphids (Dixon et al. 1993). This is thought to be due to energy limitation, as allocation of resources to the muscles reduces the availability of the same for reproductive functions. On the other hand, several investigators have reported a positive correlation between dispersal and fecundity (Hanski et al. 2006, reviewed in Rankin and Burchsted 1992), which is possible if the dispersers are also the physically superior organisms within the population who abound in resources (Bonte and de la Peña 2009). However, our results differed from both these expectations, and there was no significant difference between the fecundity of the VB and the VBC populations in either early or late life (Fig. 3B), which is consistent with an earlier dispersal evolution study on spotted-mites (Fronhofer et al. 2014). Similarly, our results did not agree with expectations from the literature in terms of longevity. In Glanville fritillary butterflies, higher dispersal and mobility correlate strongly with higher flight metabolic performance (Hanski et al. 2004). Since a strong negative correlation between life-span and metabolic rate has been reported across various taxa (reviewed in Rattan 2008), dispersers are expected to have a shorter life-span, which was actually observed in a previous study on tropical butterflies (Tufto et al. 2012). However, we did not find any significant difference in longevity between the dispersal-selected and the control populations, a result which is consistent over all four populations and both the sexes (Fig. 3C). Although these lack of changes in fecundity and longevity are puzzling at the first glance, we note here that even after 67 generations (when the last set of assays reported in this study were performed), the VB populations were still responding to selection for dispersal ability. This was evident from the fact that the distance between the source and the destination were still being increased every few generations during selection. Therefore, it is possible that the selected flies had still not evolved sufficiently high dispersal ability to face statistically-significant costs in terms of fecundity and longevity. This is consistent with the observation that the actual values of average longevity of both males and females of the VBs were less than those of the VBCs (Fig 3C).

Thus we find that the individuals of the selected lines neither had greater body resources nor a reduced level of other energy-intensive traits like fecundity and life-span. Then how did they maintain an elevated level of dispersal and activity compared to the controls? To investigate this question, we next compared the metabolomes of the selected and the control flies.

### 4.6 Selected flies have elevated levels of cellular respiration

There was a clear difference between the overall metabolite profiles of the VB and VBC flies (Fig. S3 and S4), and the levels of 14 metabolites were significantly different between these two populations (Table 1). Most notably, the level of glucose, which is the primary proximate source of energy in the cell through the process of cellular respiration, was significantly higher for VBs compared with VBCs. Moreover, the VBs had greater levels of citric acid, nicotinamide adenine dinucleotide (NAD) and adenosine monophosphate (AMP), all of which are critically associated with cellular respiration (Fig. 4). Finally, the VBs also had significantly greater amounts of lactic acid. It is known that when the demand for energy is more than what cellular respiration can generate (e.g. during intense muscular activity), glucose undergoes anaerobic oxidation via lactic acid fermentation to produce ATP. True to this observation, the ATP levels in VBs were significantly higher than in the VBCs (Table 1). However, the AMP: ATP ratio, which is an indicator of levels of cellular energy crunch (Hardie and Hawley 2001), was much higher for VBs. This suggests that, in spite of the greater levels of ATP, the VBs are in a more energy-depleted state than the VBCs (Table 1). Taken together, these results suggest that the VBs have elevated levels of both aerobic and anaerobic cellular respiration, which is consistent with the fact that they disperse to longer distances (Tung et al. 2018) and have greater locomotor activity (Fig. 1A, 1C and 1E).

**Figure 4.**
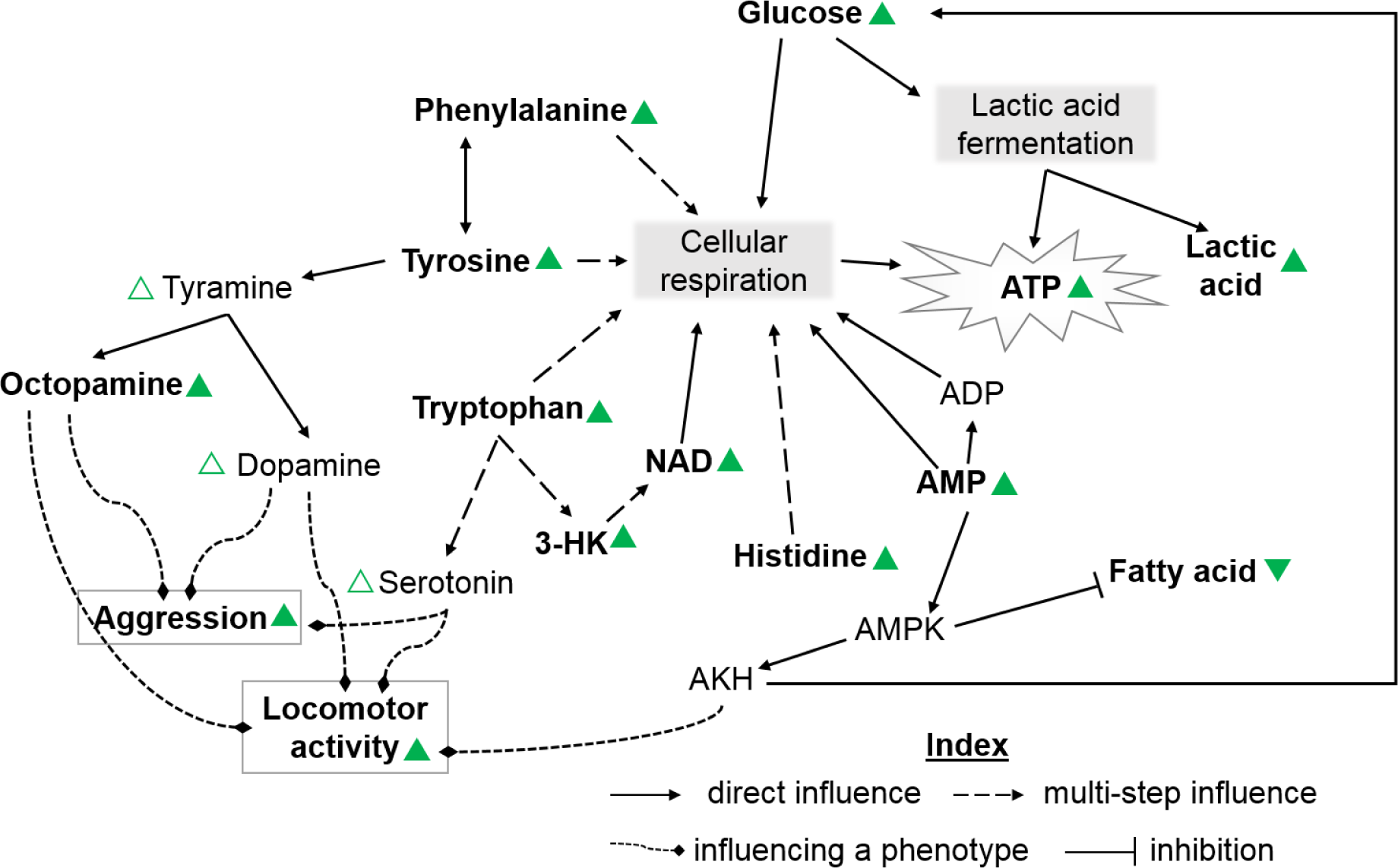
Alterations in metabolic pathways. Schematic diagram of the interactions between the metabolites that were found to be significantly changed in the VBs in the NMR data. Upright and inverted triangles adjacent to a metabolite denote whether its level had increased or decreased, respectively. Filled and open triangles represent statistically significant (see Table 1 for values) and non-significant changes, respectively.

### 4.7 Selected flies have elevated levels of octopamine and precursors for other neurotransmitters

The VBs also had increased levels of four amino acids, namely histidine, phenylalanine, tyrosine and tryptophan (Table 1). The metabolic break-down products of all these amino acids form intermediates of citric acid cycle or their precursors, and therefore play a role in energy production (Voet and Voet 2011). More interestingly, phenylalanine and tyrosine act as precursors for octopamine and dopamine (Brandau and Axelrod 1972), while tryptophan is a precursor for serotonin (Stone and Darlington 2002). In *Drosophila*, increased levels of octopamine not only enhances aggression (Zhou et al. 2008) but also leads to greater activity (Yellman et al. 1997). This was consistent with the observation that octopamine levels had significantly increased in VBs (Table 1). Similarly, serotonin levels are also known to be positively correlated with activity (Yellman et al. 1997) and aggression (Dierick and Greenspan 2007). Dopamine can elevate the activity level in flies (Yellman et al. 1997), although its relationship with aggression is not as straightforward as for the other molecules (Alekseyenko et al. 2010). Combining these evidences with the fact that the VBs are more active (Fig. 1A, 1C and 1E) and aggressive (Fig. 2B), there is a strong indication that the levels of serotonin and dopamine have also increased in the course of dispersal evolution. We did detect an increase in the levels of both these neurotransmitters with large (*d*= 1.16) and medium (*d*= 0.73) effect sizes respectively, although the increase was not statistically significant after Bonferroni correction.

It should be noted here that apart from the three neurotransmitters discussed above, there are many more molecules which can also potentially modulate fly behaviour. For example, it is known that increased levels of insulin (Belgacem and Martin 2006; Luo et al. 2014) and tachykinin (Asahina et al. 2014) can reduce either aggression or activity or both. Unfortunately, although a comprehensive investigation of the changes in the levels of the various neurotransmitters in these flies would be of immense interest, it is also outside the scope of the present study.

### 4.8 Selected flies have reduced levels of fatty acids

The end-product of the main route of tryptophan metabolism is nicotinamide (Stone and Darlington 2002), which subsequently produces NAD, a key element of cellular respiration (Khan et al. 2007). One of the main intermediates of the tryptophan-NAD pathway, 3-hydroxykynurenine (3-HK), is also found to be significantly higher in VBs. 3-HK is associated with free-radical generation and neural degeneration in flies (Savvateeva et al. 2000), which is consistent with the slightly lower (although not significant) longevity of the VB flies (Fig. 3C). Histidine, another amino acid with elevated levels in the VBs, is known to be coregulated with AMP (Rébora et al. 2005). Abundant supply of AMP and depletion of ATP (Table 1) increases the AMP:ATP ratio, which in turn is expected to activate AMP-activated protein kinase (AMPK) (Hardie and Hawley 2001) in VBs. AMPK typically functions to facilitate the depletion of fat storage (Sinnett and Brenman 2016) which is corroborated by the observation that VBs had significantly lower levels of fatty acid (Table 1). AMPK also controls the normal secretion of adipokinetic hormone (AKH) (Braco et al. 2012). AKH in turn stimulates locomotor activity and helps in maintaining a hyperglycaemic state in the body (Bharucha et al. 2008): two facts that are consistent with our observations on VBs.

Taken together, it is evident that the changes at the behavioural and life-history level are well correlated with the underlying metabolomic changes (Fig. 4).

## 5. Conclusions

Our study shows that, in terms of the relationship between dispersal and behavioural traits, there is excellent correspondence between the insights derived from association studies on field populations and experimental evolution studies. One reason for this might be that active dispersal is intimately related to locomotion which shares common control mechanisms with aggression and exploration via neurotransmitters like octopamine and serotonin. This automatically leads to the prediction that in passively-dispersing organisms, this trait-association is likely to break down. Incidentally, we also show that in terms of life-history traits, the correspondence between field and laboratory studies is poor. Finally, our study gives the first glimpses of the metabolome-level changes that accompany dispersal evolution. This is best thought of as an overview of the myriad changes that can occur when dispersal evolves, and the complex ways by which these changes can affect the various traits of the organism. Establishing the robustness of these metabolic level changes (particularly under field conditions) and connecting them to the corresponding genes is going to be one of the next big challenges in dispersal ecology.

## Acknowledgements

We thank Partha Pratim Chakraborty and Selveshwari S. for help during the experiments, Jeet Kalia for allowing us to use the Lyophilizer for NMR sample preparation; and Amitabh Joshi, Sagar Pandit and Milind Watve for helpful discussions. ST and AM were supported by Senior and Junior Research Fellowships, respectively, from the Council of Scientific and Industrial Research, Government of India. MAS and PMS were supported through the INSPIRE fellowship of Department of Science and Technology, Government of India. VRSS was supported through the GE Foundation Scholar Leaders Program. NG and KD thank the NMR Research Facility at IISER Mohali. NG was supported by an IISER Mohali institutional fellowship. This study was supported by a research grant (#EMR/2014/000476) from DST, Government of India and internal funding from IISER-Pune.

**Data Statement:** Data will be deposited in Dryad if accepted.

## Appendix S1: Detailed materials and methods

### Text S1.1 Ancestry and maintenance regime of the experimental populations

In this experiment, we used eight large (breeding size of ~2400) outbred laboratory populations of *Drosophila melanogaster*, four of which were subjected to selection for increased dispersal over 49 generations (called VB_1-4_) and the other four populations served as corresponding controls (VBC_1-4_). VB and VBC populations that share a numerical subscript (e.g. VB_1_ and VBC_1_) were related by ancestry (Tung et al. 2018), and hence were always assayed together and treated as blocks in statistical analyses. The ancestry of these flies traces back to the IV lines, which were wild-caught at South Amherst, MA, USA in 1970 (Ives 1970). These flies have been maintained ever-since at large population sizes (to ameliorate inbreeding-like effects) under laboratory conditions.

Both VBs and VBCs were maintained on a 15-day discrete generation cycle at 25°C and constant light conditions. In each generation, ~60-80 eggs were collected in clear plastic vials containing ~6 mL of standard banana-jaggery food (following Sheeba et al. 1998). For each VB and VBC population, we collected eggs in 80 and 40 vials respectively. After 12 days from the day of egg collection, the adults were collected and subjected to the selection protocol (see Text S1.2). Immediately after this, the adults of a given population were transferred to a plexi-glass cages (25 cm χ 20 cm χ15 cm) and provided with yeast supplement along with standard banana-jaggery food. After ~40 hours, eggs were collected for the next generation. The adults were discarded after oviposition, thus ensuring that individuals of two successive generations never co-exist.

### Text S1.2 Assays

Prior to any assay, both VB and VBC populations were maintained under identical rearing conditions for one generation to ameliorate the influence of phenotypic plasticity or non-genetic parental effects. The progeny of these flies were used for the assays. Moreover, for all the assays, egg density was always maintained at ~50 eggs on ~6mL food in each vial to avoid any confounding effect of larval crowding on the life-history and behavioural traits measured.

#### 1 Locomotor activity assay

After 49 generations of selection, locomotor activity of the selected and control lines were checked both in the presence and absence of food using *Drosophila* Activity Monitoring (DAM2) data collection system (Trikinetics Inc, Waltham, MA). For the activity assay in the presence of food, on the 11^th^ day from the day of egg collection, between 1830 h -1930 h, single adult male flies were aspirated into glass activity tubes of 5 mm diameter and 6.5 cm length, containing standard banana-jaggery food at one end. We preferred aspiration over CO_2_ anaesthesia to avoid any lingering effects of anaesthetization on the activity of the flies (van Dijken et al. 1977). Details of the preparation of the tubes and the cleaning of the same can be found elsewhere (Chiu et al. 2010). During the assay, the tubes (each containing a single fly) were placed in the monitoring apparatus such that two independent perpendicular IR beams passed through the centre of each glass tube. The activity for a given fly was estimated as the average number of times the fly crossed the IR beam per hour (Chiu et al. 2010). Continuous inactivity for five minutes or more was considered as sleep/rest (Hendricks et al. 2000; Chiu et al. 2010). For each VB/VBC population, we measured the activity of only male flies as in females laying of eggs on the tubes could affect the accurate measurement of locomotor activity (Chiu et al. 2010). The selected populations were always assayed along with their matched control populations (i.e. VB_1_ was assayed with VBC_1_ and so on) and there were 30-32 replicates for each population. Activity data were collected for 30 hours and were divided into two parts - i) first 6 hours and ii) next 24 hours. The first set captured the activity-rest pattern immediately after introduction of the flies in the tube, while the next set measured the steady state activity-rest pattern, after 6 hours of acclimatization, for a complete 24-hour cycle. For the entire duration of recording, the monitors containing the activity tubes were kept undisturbed inside an incubator maintained at 25 °C and constant light.

Locomotor activity assay in the absence of food was similar to the one mentioned above except that no food was provided and both ends of the activity tubes were secured with clean, dry cotton plugs. The setup for this assay was done on the 12^th^ day from the day of egg collection between 1200 h-1300 h which roughly corresponds to the time at which selection was imposed during the regular maintenance of VBs. Moreover, in the no-food case, locomotor activity was recorded only for 6 hours from the time of setup as, after this period, the flies become stressed and slowly start dying due to desiccation. 30-32 flies were assayed for each of the VB and VBC populations.

For each of these datasets, average number of activity counts per hour was calculated as a measure of the activity level of the flies and the fraction of time the flies remained inactive was computed as an indicator of the sleeping/resting duration. Note that activity and rest are defined on different blocks of time, and therefore are independent of each other. For example, a given fly can potentially rest/sleep for longer period and be more active by running back-and-forth excessively in the tube, while it is awake/not resting. In order to assess the quality of rest/sleep, for the 24-hour dataset, we computed the average length of uninterrupted rest/sleep bout and duration of the longest rest/sleep bout for each of the flies.

#### 2 Exploration assay

This assay was performed after 53 generations of selection. For each VB or VBC population, we used 10 replicate vials each containing around 50 eggs / 6ml of banana-jaggery food. The assay was performed on the 12^th^ day after egg collection. The male flies were aspirated from the egg-collection vials and introduced into the experimental arena (modified from Soibam et al. 2012) where their activity was recorded using a video camera. The experimental arena was made of a clear polycarbonate petri dish lid with an inner diameter of 10 cm. A small hole was drilled into the centre of the lid to introduce flies into the setup. In general, the flies prefer to walk along the boundary of the arena (i.e. side-wall of the lid), avoiding the inner zone (Soibam et al. 2012). Thus, movements away from the boundary indicates exploratory tendency. To score this behaviour, we placed the petri-plate lid on top of a blank white sheet of paper which contained the traces of two concentric circles. The outer circle was of the same diameter as the petri dish lid while the inner circle divided the total area of the lid into two zones: the outer containing one third of the total area while the inner one enclosing two-thirds of the area (Liu et al. 2007) of the lid. 32 replicates were assayed for each sex of all the VB and VBC populations. After the flies were introduced into the arena, they were given one minute to acclimatize to the new environment. They were then observed for the next 10 minutes and the number of times they entered the inner zone, considered as the number of exploratory trips, was recorded.

#### 3 Male-male aggression assay

The aggression assay was performed after 52 generations of selection. For this assay, the flies were reared under low levels of larval crowding (~50 eggs in 6-8ml banana-jaggery food). Freshly eclosed males were collected and reared in social isolation (i.e. one male per vial) till the day of assay (following Yurkovic et al. 2006). We used 6 wells of a twelve-well culture plate (Corning®, NY, USA) as the assay apparatus, where each well served as the enclosure for one replicate of the aggression assay. A small plastic cup containing regular banana-jaggery medium was affixed at the centre of each well. A freshly decapitated female was stuck to the middle of the food cup using yeast paste. The food and the female served as defendable resources and potential reasons for conflict. Following an earlier protocol, VB and VBC males were colour-coated with daylight fluorescent pigments (DayGlo) for easy identification (Dickens and Brant 2014). On the 12^th^ day from egg collection, two males (one VB and one VBC) were introduced into the setup and their interaction was recorded for 45 minutes using a video camera. 30 such replicates were assayed for each of the four populations of VBs and the corresponding VBCs. Individual wells were visually isolated from each other using cotton to ensure no visual cues were being exchanged between replicates. Uniform lighting and constant temperature (25°C) were maintained.

The scoring for aggression was done after an initial five-minute acclimation period. For each of the replicates, the number of successful chase-aways from the food cup was recorded. A successful chase-away is defined as one in which one male completely chases the other male away from the top surface of the food cup (Yurkovic et al. 2006). Earlier studies have shown that in *Drosophila* a male that manages to complete three consecutive successful chase-aways usually manages to successfully chase away the other male in all future encounters (Yurkovic et al. 2006). Therefore, we used this criterion to identify the winner of each fight from all the blocks.

#### 4 Body size assay

Dry body weight of the adults was measured as a proxy for body size after 49 generations of selection. For a given population, ~50 eggs were introduced into food vials containing ~6-8 mL of standard banana-jaggery medium. After 12 days from the day of egg collection, the adult flies were collected, sorted by sex, killed by flash freezing and stored at -80 °C till weighing. The flies were then dried at 60°C for 72 hours and (after thawing to room temperature) weighed to the nearest 0.1 mg in batches of 20 males or 20 females. Five batches of males and females were weighed for each of the four VB and VBC populations.

#### 5 Female fecundity assay

Female fecundity of the selected (i.e., VB_1-4_) and the control (i.e. VBC_1-4_) populations were assayed after 53 generations of selection. Female fecundity was assayed both during early and late life. For early life fecundity, we used 15-day (post egg collection) old females, which is the same age at which the eggs are collected for the selection lines. Late life fecundity was measured on 33-day (post egg collection) old females. This is because, in *Drosophila*, it is known that during this time female fecundity reduces substantially due to aging but does not plateau out (Hanson and Ferris 1929). Till the respective day of assay the flies were maintained in mixed-sex groups of ~50 individuals on ~6mL of standard banana-jaggery food. Unlike normal maintenance regime, the adult flies were not provided excess yeast paste to boost their fecundity (Mueller and Huynh 1994). On the day of assay, flies were anaesthetized under mild CO2 and one male and one female each were introduced into a 50 mL centrifuge tube containing a food cup. The tube had provision for aeration and the food in the food cup provided a surface for laying eggs. 40 such replicate setups were made for each VB_*i*_ and VBC_*i*_ (where *i* ∈ 1-4) population. The setups were left undisturbed for 12 hours in a well-lit environment maintained at 25°C and ambient humidity. At the end of 12 hours, the flies were discarded and the eggs laid on the food were counted under a stereo microscope. Since fecundity is largely determined by the body size of the females (Honek 1993) which is in turn critically dependent on larval density (Prout and McChesney 1985), we maintained a constant egg density (~50 eggs per vial containing ~6-8 mL of standard banana-jaggery food), while collecting eggs for generating the flies for this assay.

#### 6 Longevity assay

Longevity assay was performed after 51 generations of selection in a constantly lit environment maintained at 25°C. We initiated the assay by introducing 10 freshly eclosed, unmated individuals of the same sex into a food vial containing ~6 mL standard banana-jaggery food. 10 such replicate setups were prepared for males and females separately from each of the VB/VBC populations. Thus in total, we measured the life-span of 1600 flies in this assay. The alive flies were counted daily at a particular time (arbitrarily set at 1500 h) and every alternate day, they were transferred into fresh food vials, till the last individual died.

#### 7 Metabolomic study using NMR spectroscopy Sample preparation

After 67 generations of selection, NMR spectroscopy was performed on one block of selected-control populations (VB_4_-VBC_4_). After rearing under common conditions for one generation, eggs were collected (~50 eggs on ~6 mL food) to generate the adult flies for studying the metabolomics of the two kind of populations. On the 12^th^ day after egg collection, for each VB/VBC population, we prepared 11 replicates, each containing 30 males and processed them using established protocols (Gogna et al. 2015). The flies were first flash-frozen in liquid nitrogen, transferred to pre-labelled microfuges containing 0.5 mL of 50% acetonitrile solution, homogenized using a battery run homogenizer and centrifuged at 10000 rpm for 10 min at 4°C. The supernatant was then transferred to another set of pre-labelled microfuge tubes, lyophilized and stored at -80°C, to be used for NMR experiments. Prior to the NMR experiments, the samples were rehydrated in 500 ml of 50 mM phosphate buffer prepared using D_2_O (pH 7.4), containing 1 mg/ ml of 3-(trimethylsilyl)-propionic acid-D4, sodium salt (TMSP) as a chemical shift reference and transferred to 5mm NMR tubes.

### NMR spectroscopy

NMR spectra were recorded on a Bruker Biospin 600 Avance-III spectrometer operating at a ^1^H frequency of 600.13 MHz at 300 K using a 5 mm QXI probe. Gradient shimming was performed prior to signal acquisition to optimize magnetic field homogeneity. 1D ^1^H NMR spectra were acquired using the water suppressed Car-Purcell-Meiboom-Gill (CPMG) spin-echo pulse sequence optimized with a spin -echo delay *t* of 300 ms and n= 400 and a total spin-spin relaxation delay (2nt) time of 240 ms to achieve attenuation of fast-relaxing broad signals from larger molecules. The proton spectra were collected with a 90-degree pulse width of 9.15 ms, a relaxation delay of 2 s, 16 scans, 16 K data points and a spectral width of 7211.54 Hz. Data were zero-filled by a factor of 2 and the FIDs were multiplied by an exponential weighting function equivalent to a line broadening of 1 Hz prior to Fourier transformation. For resonance assignment and metabolite identification, two - dimensional NMR spectra were recorded, including ^1^H-^1^H correlation spectroscopy (COSY) and ^1^H-^13^C heteronuclear and homonuclear single quantum coherence spectroscopy (HSQC, HMQC). 2D ^1^H-^13^C HMQC and HSQC spectra were obtained with a spectral width of 12 ppm and 200 ppm in the proton and carbon dimensions respectively, 1 K data points, 32 scans, 256 t1 increments and a recycle delay of 1.5 s. The COSY spectra were acquired with a spectral width of 12 ppm in both dimensions, 2 K data points, 32 scans and 128 t_1_ increments. Metabolite fingerprinting for the *Drosophila* NMR spectra was done by checking identified metabolite peaks with standard NMR metabolite data deposited in databases such as MMCD (http://mmcd.nmrfam.wise.edu) and BMRB (http://www.bmrb.wise.edu). The NMR chemical shift assignments of several significant metabolites were further confirmed by recording the NMR spectra of pure compounds. For analysis of metabolites, single peak integrals for individual metabolites were chosen with minimal overlaps with peaks from other compounds.

### Data Analysis

Multivariate statistical analysis was performed using SIMCA14.0 software (Umetrics, Umea, Sweden). Prior to analysis, all the spectra were converted into the ASCII format and imported into MATLAB for alignment using the Icoshift algorithm (Savorani et al. 2010). Spectral regions between 4.6 and 4.8 ppm were excluded from the analysis, to prevent errors due to any residual peak from the suppressed water signal. Data were normalized to the total area to compensate for possible differences in signal-to-noise ratios between spectra and to prevent separation due to variations in the amounts of sample.

After importing the data into SIMCA, the data was Pareto-scaled and first analysed using the unsupervised pattern recognition method of principal component analysis (PCA), which helped to remove outliers, defined in the data as observations located outside the 95% confidence region of the Hotelling’s T^2^ ellipses in the PCA score plots. Such outliers were excluded from further analysis. PCA was followed by the supervised pattern recognition method of orthogonal projections to latent structure-discriminant analysis (OPLS-DA), which maximizes the class discrimination. The OPLS-DA scores and loadings plots were used to identify the metabolites responsible for separating VB and VBC flies. The quality of the model was described by *RX^2^* and *Q^2^* values, explaining the variance explained (indicating goodness of fit) and variance predicted by the model (predictability) respectively. The significance test of the model was performed using CV-ANOVA (cross-validated ANOVA) in the SIMCA software, where a*p* -value of 0.01 was considered to be statistically significant to validate the OPLS-DA model. Permutation analysis was also performed on the best model using 1000 permutation tests with a threshold *p*-value of 0.05. t-tests coupled with Bonferroni corrections (to limit the family-wise error rate to 0.05) were performed to check for statistical significance of the differences in the metabolite levels between the VB and VBC flies.

**Figure S1.**
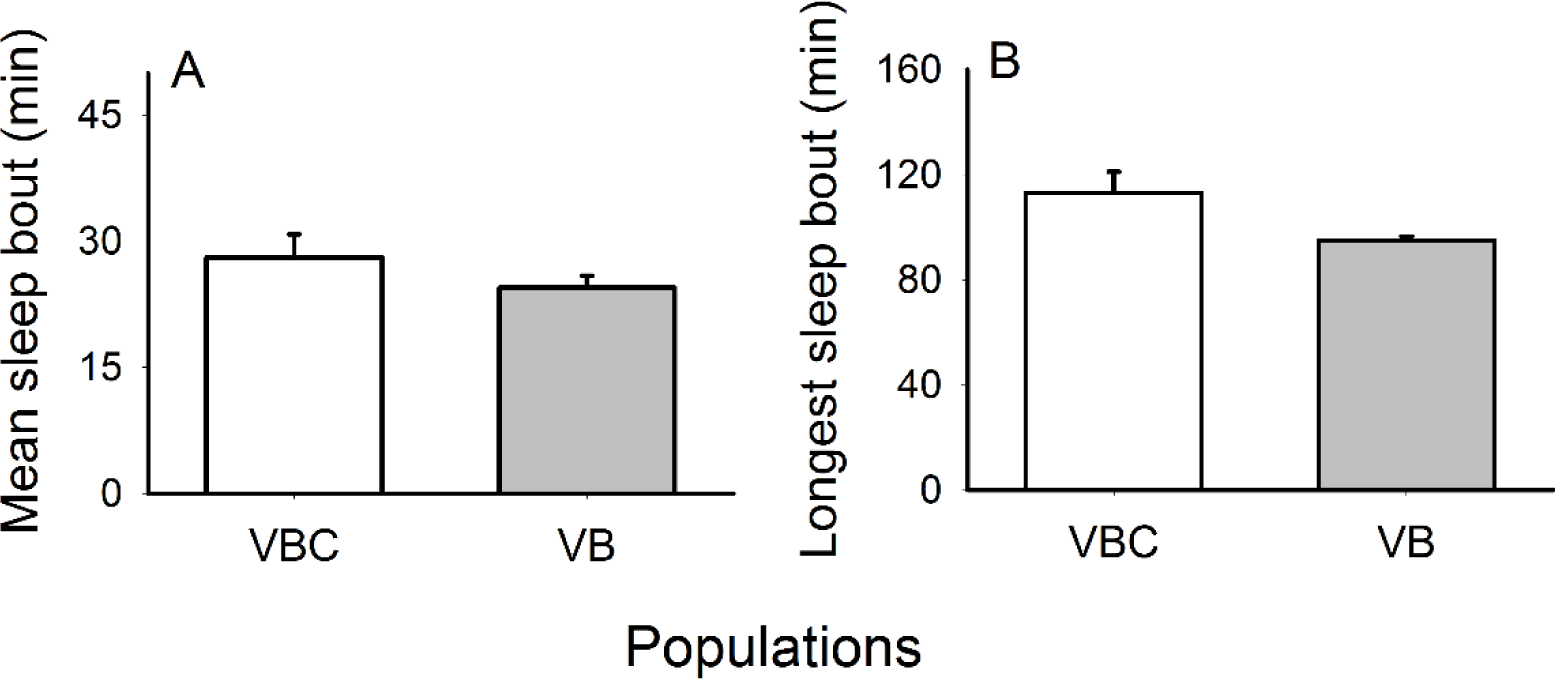
Average and maximum sleep bout over 24 hours post acclimatization. Over 24 hours in presence of food, (**A**) average sleep bout and (**B**) maximum sleep bout, are similar for both VBs and VBCs. The error bars represent standard errors around the mean (SEM).

**Figure S2.**
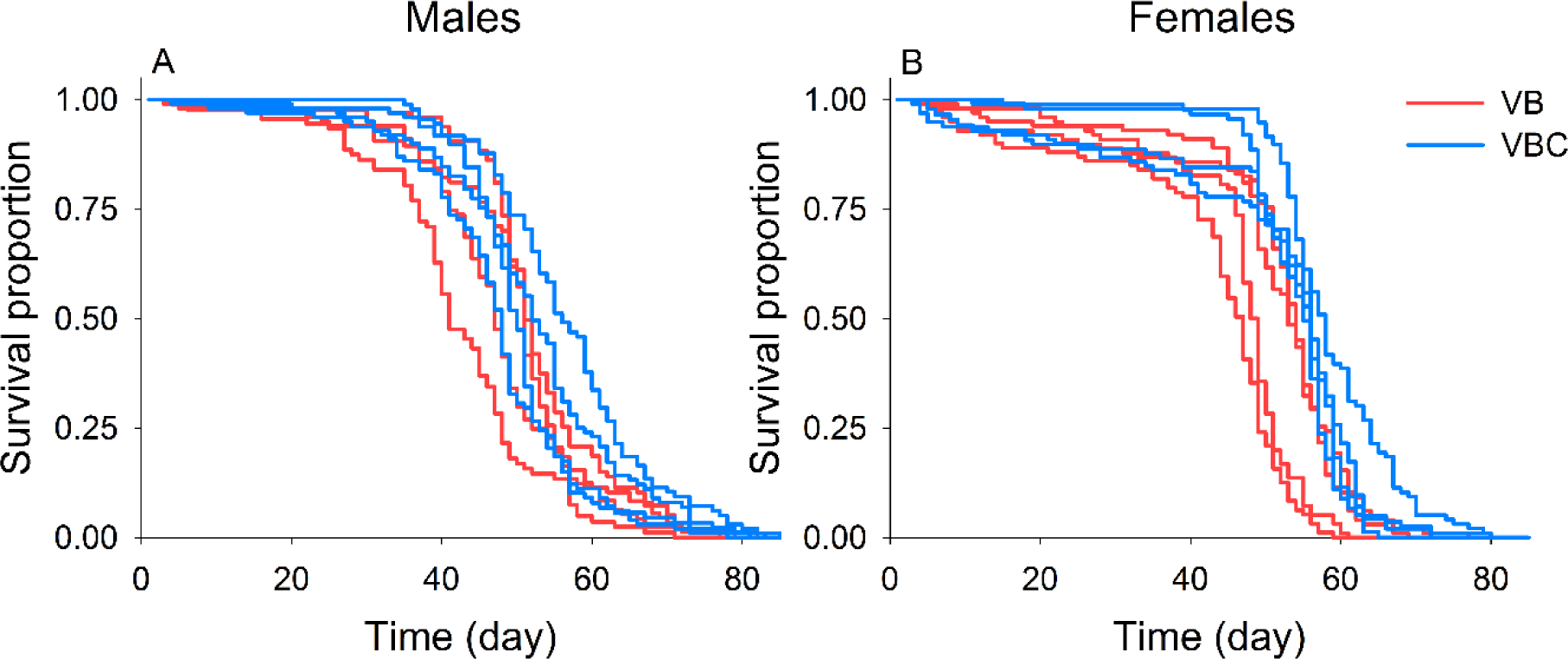
Survival curves for VB and VBC populations. The plots depict the probability of survival as a function of time based on Kaplan-Meier estimates, for (A) males and (B) females of all four VB (red) and VBC (blue) populations. Although there is a trend, survivorship curves of the two groups do not differ significantly, when compared using Cox proportional hazards model including the random factor (using ‘coxme’ in R package ‘coxme’) as well as excluding the random factor (using ‘coxph’ in R package ‘survival’).

**Figure S3.**
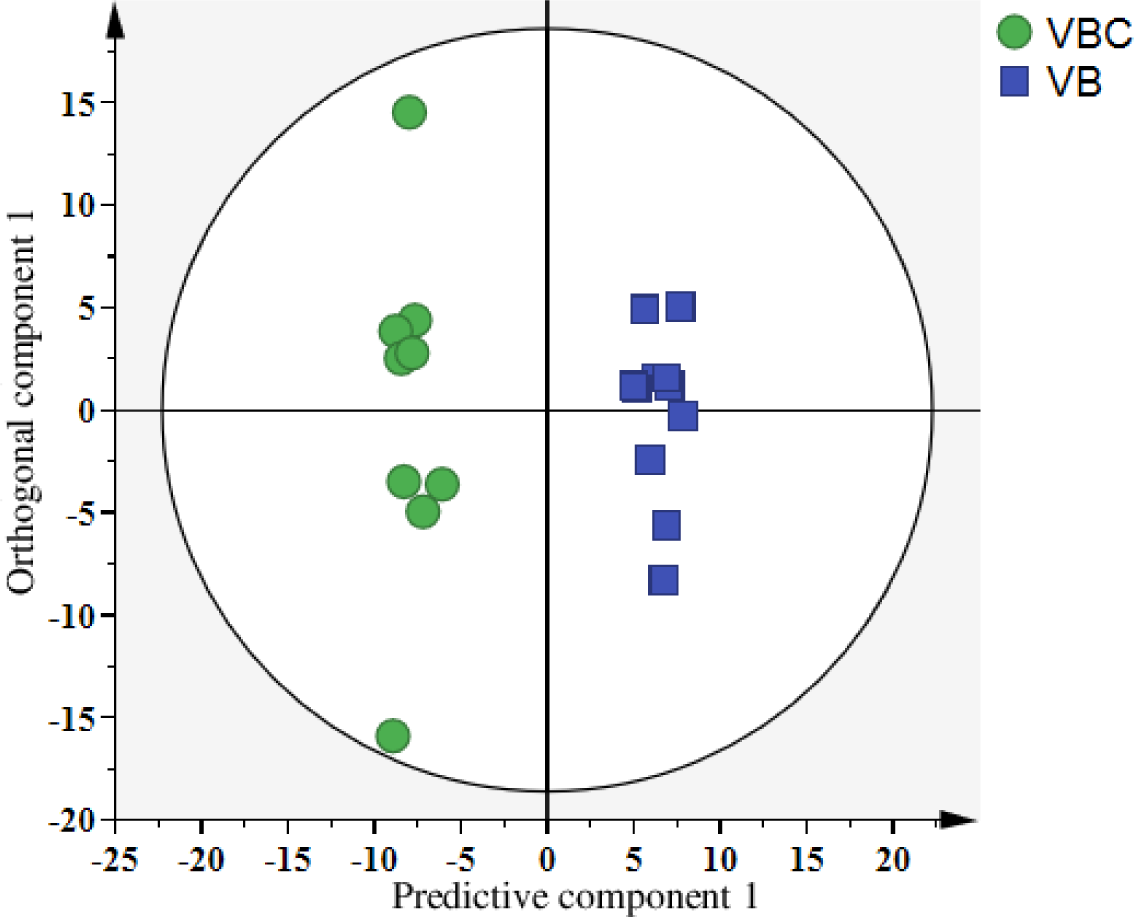
OPLS-DA score plot derived from 1D ^1^H NMR spectra of VBs and VBCs. The plot shows that all the replicates of VB and VBC population form non-overlapping clusters indicating distinct metabolomics signatures in these two groups of flies.

**Figure S4.**
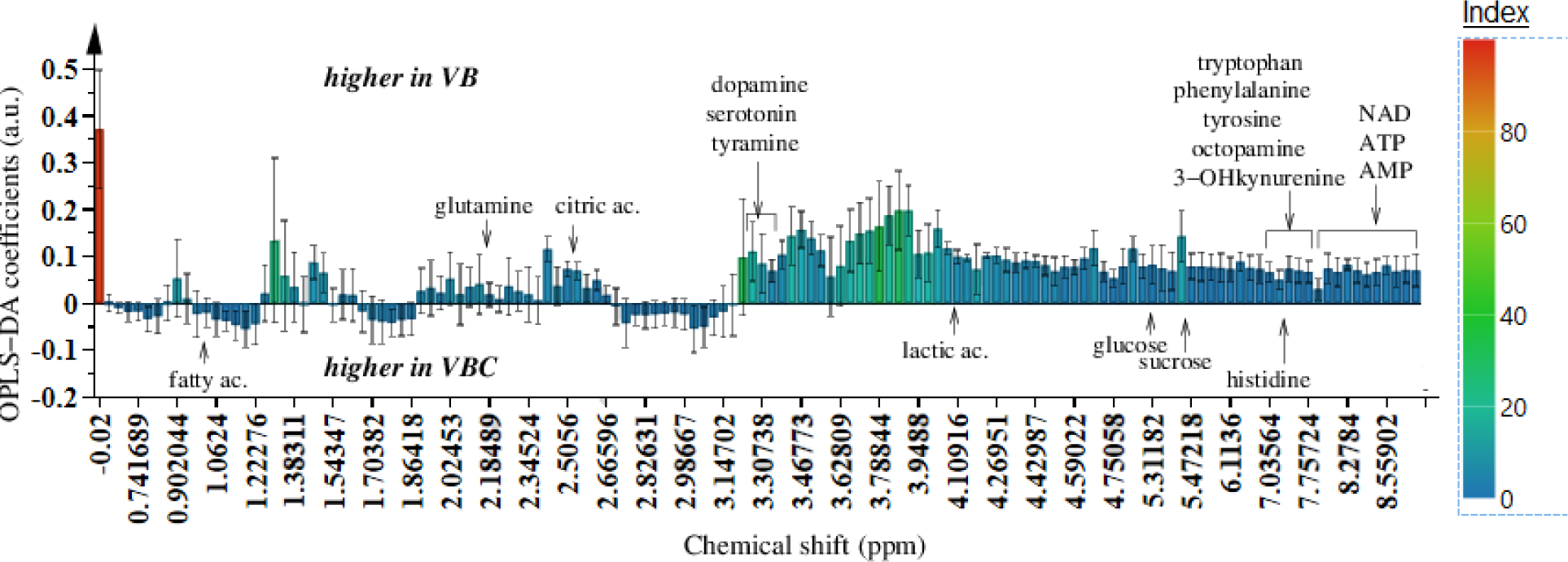
OPLS-DA loading plot obtained from the analysis of 1D ^1^H NMR spectra of VBs and VBCs. Metabolites that are indicated above and below the baseline are present in higher quantity in VB and VBC flies respectively.

